# The view of microbes as energy converters illustrates the trade-off between growth rate and yield

**DOI:** 10.1101/2021.04.16.440103

**Authors:** St. Elmo Wilken, Victor Vera Frazão, Nima P. Saadat, Oliver Ebenhöh

## Abstract

The application of thermodynamics to microbial growth has a long tradition that originated in the middle of the 20^th^ century. This approach reflects the view that self-replication is a thermodynamic process that is not fundamentally different from mechanical thermodynamics. The key distinction is that a free energy gradient is not converted into mechanical (or any other form of) energy, but rather into new biomass. As such, microbes can be viewed as energy converters that convert a part of the energy contained in environmental nutrients into chemical energy that drives self-replication. Before the advent of high-throughput sequencing technologies, only the most central metabolic pathways were known. However, precise measurement techniques allowed for the quantification of exchanged extracellular nutrients and heat of growing microbes with their environment. These data, together with the absence of knowledge of metabolic details, drove the development of so-called black box models, which only consider the observable interactions of a cell with its environment and neglect all details of how exactly inputs are converted into outputs. Now, genome sequencing and genome-scale metabolic models provide us with unprecedented detail about metabolic processes inside the cell. However, the derived modelling approaches make surprisingly little use of thermodynamic concepts. Here, we review classical black box models and modern approaches that integrate thermodynamics into genome-scale metabolic models. We also illustrate how the description of microbial growth as an energy converter can help to understand and quantify the trade-off between microbial growth rate and yield.

**Perspective:**

1. Microbial growth is the foundation of many biotechnological applications. The key to optimizing microbial growth lies in thermodynamics, similar to how classical thermodynamics helped optimize steam engines in the 19^th^ century.
2. Genome-scale metabolic models have become widely available, and are used to predict microbial growth. These predictions often fail because these models do not distinguish between growth rate and yield.
3. Classical black box models present a sound thermodynamic theory, by viewing microbes as energy converters. Incorporating such concepts into genome-scale metabolic models has the promise to advance our fundamental understanding of microbial growth, and thus to improve the predictive power of these models.

## 1 Introduction

Thermodynamics has its origins in the optimization of steam engines in the 19^th^ century. During this era, two of the fundamental laws of thermodynamics were postulated: the first law holds that energy is conserved in an isolated system; and the second law holds that no process is possible that only transfers heat from a lower temperature to a higher temperature^1^ [1]. While the resultant theory mostly dealt with systems at equilibrium, in the middle of the 20^th^ century strides were made to extend it to irreversible, non-equilibrium regimes, allowing for the direct analysis of less idealized systems [2, 3]. Today, thermodynamics is integral to understanding efficiency limitations of natural and engineered systems, and has been widely applied in biochemistry and biotechnology [4, 5, 6, 7, 8].

Efficiency considerations guide researchers in their quest to rationalize the design of living systems in light of evolutionary optimization. The importance of understanding efficiency limitations is further stressed by the fact that the economic competitiveness of bio-processes is often seen as a major hurdle in the development of a sustainable bio-economy [9]. Because many biotechnological processes are based on exploiting microbes as bio-based factories [10, 11, 12], the continued development and application of thermodynamic concepts and tools to study and optimize microbial growth processes is necessary. Heterotrophs obtain energy by metabolizing higher energy nutrients into lower energy components through a process known as catabolism. Since a part of this energy gradient is used to build new biomass (anabolism), heterotrophic growth can be seen as a thermodynamic energy converter [13], where ATP is the central coupling metabolite, which facilitates the conversion of energy released in catabolic processes to drive biomass formation. This thermodynamic view of microbial growth opens the door to a rich array of tools that can be used to analyze these systems [14].

A complete understanding and quantitative description of heterotrophic growth will depend on the knowledge of the detailed mechanisms of how this energy converter functions. Historically, a black box description was used to investigate the overall conversion [15], but this approach did not reveal how the coupling is performed in detail. For example, it is unknown how many molecules of ATP are being produced per carbon consumed in catabolism, and how many are needed to incorporate one carbon into biomass in anabolism. Genome-scale metabolic models (GEMs) reveal a view into the black box by mapping pathways and reactions to specific genes, and are increasingly used in systems biology applications [16], but often do not adequately reflect thermodynamic constraints [17]. It is apparent that a mechanistically sound description of the microbial energy conversion process must obey the laws of thermodynamics. Incorporating concepts from thermodynamic black box models into GEMs therefore seems to be a promising way forward to develop a thermodynamically consistent, and mechanistically detailed, description of metabolism.

In this review, we first discuss the basis for all further thermodynamic calculations used to describe biological systems (Section 2). This basis is given by the concept of Gibbs free energies of reactions. We discuss its importance, its connection to biochemistry, existing methods to calculate it, and how it can be applied to glean insight into single reactions and complete metabolic pathways. We then review the black box approaches used to investigate microbial thermodynamics (Section 3), before we review and discuss more recent techniques that make use of GEMs for this purpose (Section 4). In Section 5 we discuss how these approaches may be connected. Viewing microbial growth as a thermodynamic energy converter provides access to key quantities, such as thermodynamic efficiency. This, in turn, immediately raises fundamental questions regarding the trade-off between growth rate (kinetic efficiency) and yield (stoichiometric efficiency). This trade-off is a recurring theme in this review, as it is encountered on various levels in different models describing microbial growth as a thermodynamic energy conversion process.

## 2 Gibbs free energies - the universal driving force of reactions and pathways

The laws of thermodynamics can be used to deduce properties of reactions. Typically, chemical reactions take place at constant temperature and pressure, leading to the use of Gibbs free energy, *G*, as the primary thermodynamic potential of interest. As a consequence of the second law of thermodynamics, the change in Gibbs free energy of a spontaneous reaction, Δ_r_*G*, can only be negative, constraining the direction of net flux for a given reaction. This potential difference can also be used to find the equilibrium conditions of a reaction through the relation Δ_r_*G*^*∘*^ = −*RT* ln(*K*), where *K* is the equilibrium reaction quotient, *T* the thermodynamic temperature, and Δ_r_*G*^*∘*^ the standard^2^ Gibbs free energy change of the reaction. Standard Gibbs free energies can be found in thermodynamic databases [1].

The practical importance of Gibbs free energy has led to the compilation of numerous databases storing thermodynamic data for biologically relevant species and reactions. In particular, two current examples of such databases include the Thermodynamics of Enzyme-Catalyzed Reactions Database [18], and eQuilibrator [19]. The latter database offers the highest coverage of biochemical reactions to date [20], while being thermodynamically consistent [21]. Importantly, these databases deviate from conventional chemical reaction thermodynamics by their use of *apparent* equilibrium constants, *K*^*′*^, as well as *apparent* standard Gibbs free energies of formation, Δ_f_*G*^*′∘*^, and reaction, Δ_r_*G*^*′∘*^, to describe the thermodynamic potential of biochemical reactions. In essence, the qualifier “apparent” denotes that pH and ionic strength are held constant when calculating thermodynamic potentials. This deviation is necessitated by practical challenges associated with biochemistry, e.g. measurements of enzymatic reactions are typically made in buffered solutions where *H*^+^ is not conserved, and protonation states of metabolites are difficult to measure independently [22]. The most important consequence of this is that chemical and biochemical thermodynamic databases are not directly compatible with each other, however, the fundamentally important relations between *G* and properties of interest remain unchanged, as shown in equations (1) – (3), where *Q*^*′*^ is the apparent reaction quotient, *v*_*i*_ is the stoichiometric coefficient associated with species *i* in a reaction with *N*^*′*^ different chemical species (excluding protons).

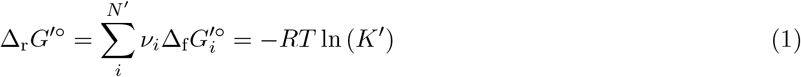

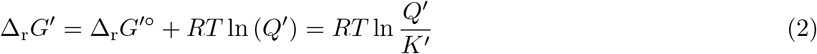

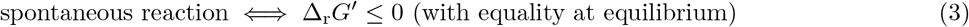

Gibbs free energy can be used to identify energetic bottlenecks in native and engineered microbial metabolism. However, the use of thermodynamic considerations in flux analysis has not been without controversy. The primary point of contention is the relationship between flux (or rate) and thermodynamic driving force. The sequence of correspondence [23, 24, 25, 26, 27] highlights the common misconception that flux is independent of the thermodynamic driving force. In contrast, under certain circumstances the flux, *J*, through an enzymatic reaction *is* proportional to the associated thermodynamic driving force (−Δ_r_*G*). The flux through a reaction can be decomposed into kinetic (*F*_K_) and thermodynamic (*F*_T_) factors, i.e. *J* = *F*_K_*F*_T_ [28, 29, 30, 31, 32, 33]. The thermodynamic factor can be written in the form 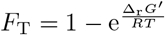 [34, 33]. If the thermodynamic driving force of a reaction is large (Δ_*r*_*G ≪* 0) the thermodynamic component approaches unity, approximately reducing the rate expression to the kinetic component only. It should be noted that even in this case the kinetic parameters of an enzymatic reaction are constrained by thermodynamics through the Haldane relation [35]. However, in the opposite case, *F*_*T*_ ≈ −Δ_r_*G* and the flux becomes proportional to the driving force. This is often observed in anaerobic environments, where the thermodynamic component plays a much larger role in constraining microbial growth [36].

Beyond describing the favourability of a single reaction, thermodynamic considerations can also be used to interrogate the interplay between rate and yield in microbial metabolism at the pathway level [37, 38]. Here, yield describes the fraction of the available free energy gradient that is used for useful chemical work, such as the production of ATP. As a consequence, a high yield means a small overall −Δ*G*^*′*^, since the energy gradient is partially conserved in ATP. In light of the foregoing, rate and yield represent a trade-off: the more efficient a pathway is in conserving energy, the lower the thermodynamic driving force becomes, resulting in a lower flux. This trade-off is the essential idea behind max-min driving force (MDF) analysis, which can be used to identify enzymes that act as thermodynamic bottlenecks in pathways. An enzyme operating close to equilibrium catalyses almost as many conversions in the reverse as in the forward direction. To overcome this inefficiency, such reactions require a high investment of cellular resources [39]. It is important to note that the concept of thermodynamic bottlenecks is quite different from that of kinetic bottlenecks (or rate-limiting steps), which exert a large control on the overall flux of the pathway [40]. MDF analysis was used to show that the Entner-Doudoroff (ED) pathway is thermodynamically more favourable than the Embden-Meyerhof-Parnas (EMP) pathway. Consequently, while the latter yields more ATP per glucose it is also more costly because it requires more enzyme. Thus, it was found that prokaryotes relying primarily on glycolysis for energy (e.g. anaerobes) tend to invest in the higher yielding, but enzymatically costlier, EMP strategy, while those that have alternative, non-glycolytic energy generating pathways (e.g. oxidative phosphorlyation in the case of aerobes) would prefer the enzymatically cheaper ED pathway [41]. Enzyme cost minimization is an extension of MDF that incorporates kinetic information to calculate the protein cost associated with a metabolic pathway [42]. This hybrid kinetic-thermodynamic approach was used to show that the ED and EMP pathways both lie close to the Pareto front of ATP yield vs. enzyme cost [43], confirming earlier studies, which rationalised the design of ATP-producing pathways by assuming that the naturally evolved pathways are close to optimal [44, 45, 6].

## 3 Thermodynamic black box approaches can be used to understand microbial metabolism

Before low cost genomic sequencing was available to interrogate the full structure of microbial metabolism, the exact metabolic transformations occurring inside a cell were relatively opaque. However, precise measurements of the heat and material (substrate consumption, product generation, and biomass production) exchanged with the environment could be made. Thus, the so-called black box techniques were developed, which depended on the measurement of macrochemical equations to summarise the conversion of nutrients into metabolic products and biomass in a single chemical formula. These techniques were used to predict cellular properties important for biotechnology without needing detailed metabolic information [13]. The entropy balance of a black box model of a cell, as depicted in Fig. 1, is given by

**Figure 1:**
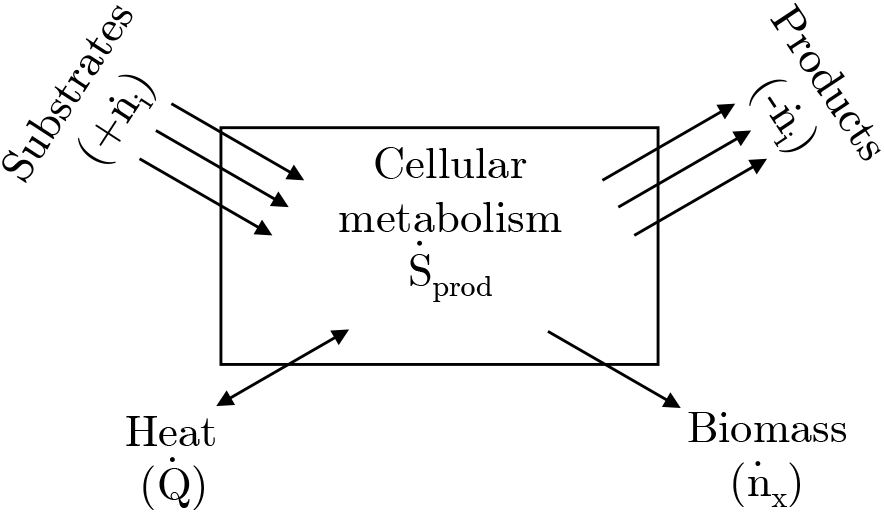
Black box approaches assume that only extracellular metabolites are measured and do not require detailed metabolic mechanisms to yield predictions. Due to the irreversible nature of cellular growth, entropy must be produced and leads to unavoidable energetic losses. The import rate of a variable is taken as positive. Based on Figure 19.10 from [46].

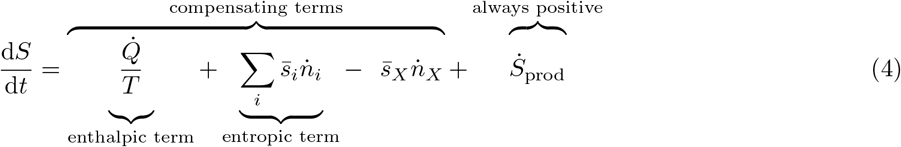

Here, 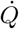 is the rate of heat exchanged between cell and environment, 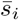 is the partial molar entropy of species *i*, 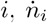 is the rate of import/export of metabolite *i*, and the subscript *X* denotes biomass. Importantly, due to the irreversibility of growth, entropy is produced by the cell 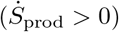, leading to unavoidable energetic losses according to the second law of thermodynamics. To avoid thermal cell death, or structural disorganization leading to cell death, a cell cannot accumulate entropy (implying 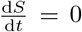). Thus, entropy production must be balanced by either heat or metabolite exchange with the environment, revealing one of the thermodynamic roles of catabolism: to export entropy in the form of heat or high entropy waste products to compensate for metabolic entropy production [46]. Interestingly, it was shown that microorganisms adopt different catabolic strategies to achieve this. Respiration is associated with enthalpic compensation, which means that entropy is exported predominantly as heat (large negative 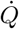), whereas fermentation is associated with entropic compensation through a high export rate (large negative 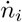) of high entropy compounds (large 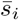). In acetotrophic methanogenesis entropy export by product formation is even so large that the overall growth is endothermic (positive 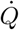). Autotrophic methanogensis, in contrast, decreases the chemical entropy in the environment, and achieves this by a very high heat production [47].

A highly simplified, but very useful and illustrative black box model, is the linear Gibbs energy converter, as illustrated in Fig. 2. In this model, catabolism drives anabolism, and these two processes are coupled through the so-called Onsager coefficients (*L*_*ij*_), as shown in Eq. (5). Here *J*_1_ and *J*_2_ are the fluxes of anabolism and catabolism, respectively; and *X*_1_ = −Δ*G*_anabolism_ and *X*_2_ = −Δ*G*_catabolism_ are the respective driving forces of these fluxes [48, 49, 50]. Note that catabolism would spontaneously proceed in the forward direction (*J*_2_ *>* 0), while anabolism, since it is endergonic, would spontaneously proceed in the negative direction (*J*_1_ *<* 0), i.e. break down biomass into its constitutive components.

**Figure 2:**
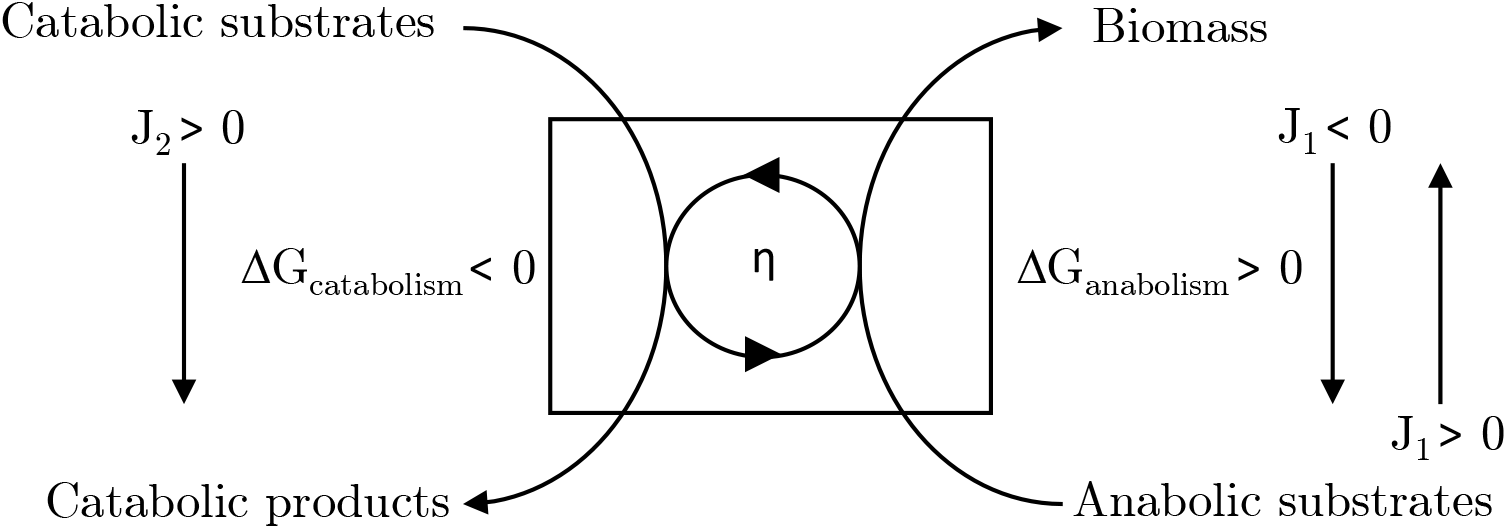
Gibbs energy converter model. Catabolism and anabolism are represented as coupled processes, where the Gibbs energy liberated by exergonic catabolic reactions drive the endergonic anabolic reactions.

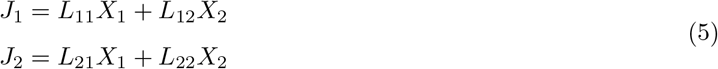

The coupling between anabolism and catabolism is effected by *L*_12_ and *L*_21_. For the purpose of studying the coupling between catabolism and anabolism, we can assume that Onsager’s reciprocal relations hold, and *L*_12_ = *L*_21_ [51]. The degree of coupling is defined as 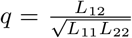. Due to the second law of thermodynamics, *L*_11_ and *L*_22_ must both be positive, but the coupling terms may be of any sign. The biologically important case occurs when they are positive, hence it can be shown that 0 *≤ q ≤* 1. In this regime catabolic free energy is channeled towards the biosynthesis of new biomass (*J*_1_ *>* 0), at the cost of catabolic flux (*J*_2_ decreases).

This simple model can be used to explain why maintenance energy is necessary in systems that are not perfectly coupled (*q <* 1). In Fig. 3a, the normalised flow ratio *j* is plotted over the normalised force ratio *x* (definitions are given in the Figure caption). Quantitatively, it can be shown that when *J*_1_ = 0 (zero net growth rate), *x* = −*q* and the catabolic flux *J*_2_ at this point is nonzero and proportional to (1 − *q*^2^) [48]. This catabolic flux represents the entropic resistance that needs to be overcome before new biomass can be produced. Maintenance energy is experimentally observed in chemostat cultures at the limit of zero growth, and, in the context of the energy converter model, corresponds to this imperfectly coupled regime.

**Figure 3:**
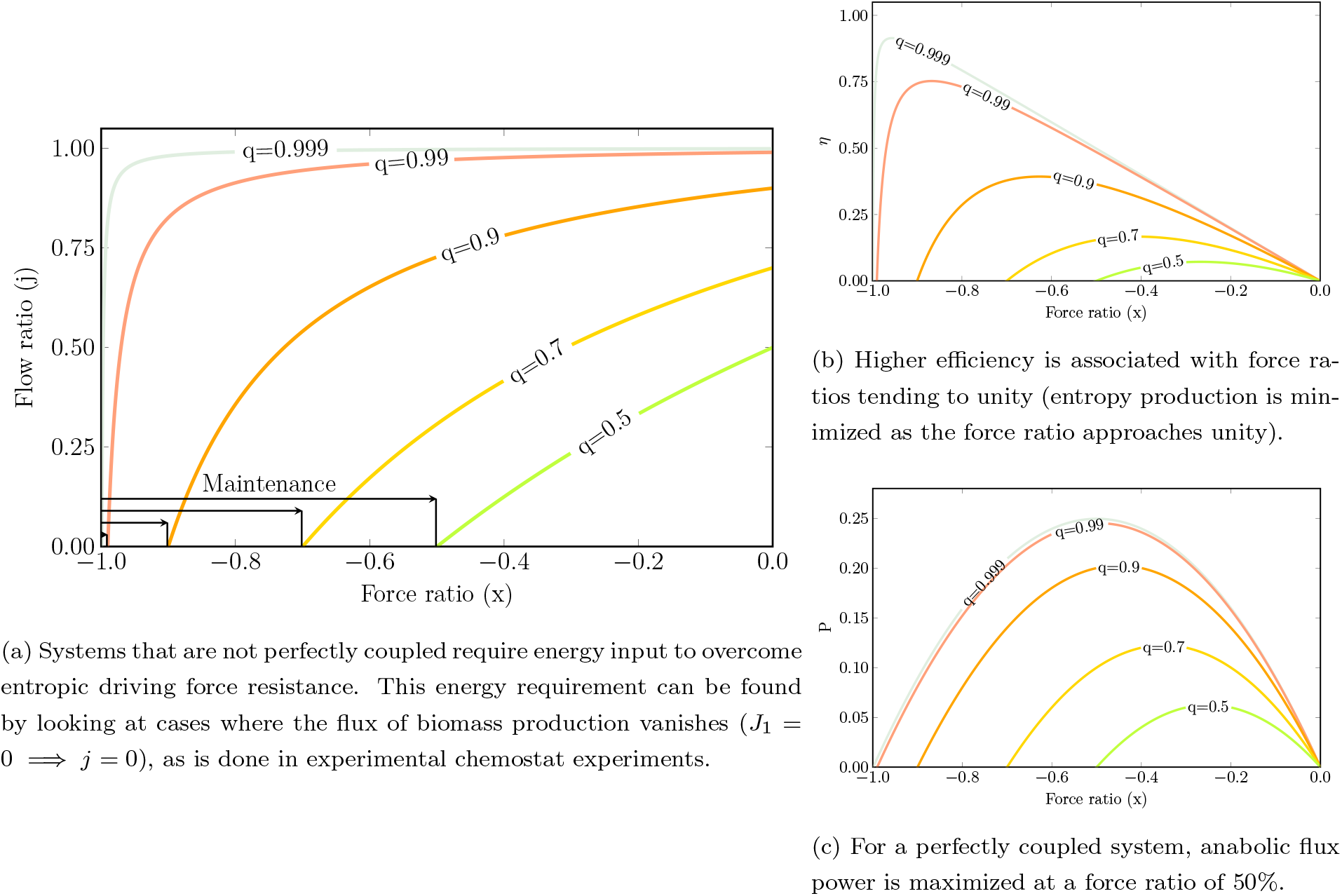
The normalized force ratio 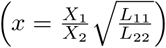 and normalized flux ratio 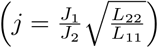 as a function of coupling strength, *q*, can be used to interrogate the predictions of the Gibbs linear energy transducer model. Positive flow ratio, *j*, indicates that biomass is being produced. Figure based on Figures 1, 2, and 5 from [48]. Code used to generate the plots are available at https://gitlab.com/qtb-hhu/thermodynamic-linear-converter-review-2021.

The efficiency, defined as 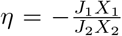, can be interpreted as the yield of the overall growth process, while the anabolic flux power (*P* = *J*_1_*X*_1_) reflects growth rate, as discussed in Section 2 for single reactions and pathways. In Figures 3b – 3c it is demonstrated that the optimum efficiency (yield) and biomass flux (rate) do not coincide, clearly indicating the existence of a trade-off between microbial growth and yield, as is often observed in nature [52].

While the linear Gibbs energy converter model is intuitively appealing, the model depends on how catabolism and anabolism are split with respect to the black box macrochemical equation, which is a non-trivial problem of definition since many catabolic reactions are actually amphibolic [13]. The problem becomes even greater if several carbon or nitrogen sources are available simultaneously. Furthermore, measuring the coupling coefficient is not straightforward. Assuming perfect coupling (*q* = 1) and maximizing anabolic flux power (*P*), the resultant predicted thermodynamic efficiency is 50%. This efficiency seems to match measured values for aerobes (*η*_observed_ *≈* 30 − 60%), but grossly overestimates anaerobic efficiencies, suggesting that the linear converter theory might be too simplistic [53].

Other approaches make direct use of the Gibbs energy balance, as shown in Eq. (6). Here *µ*_*i*_ is the chemical potential of species *i*, and 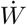 the rate of work done on the system.

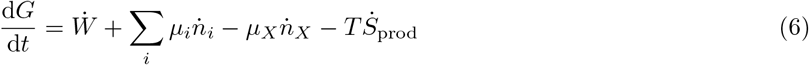

After some simplifications, it can be shown that 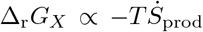, suggesting that the total Gibbs energy dissipated (Δ_r_*G*_*X*_) by a microbe is the driving force of metabolism [54, 55]. Based on this, some black box approaches have been developed that make use of Gibbs energy dissipation as a predictive tool [15]. Despite the complexity of metabolism eschewed, these techniques have been used to model biomass yield (*Y*_*X/S*_) on a variety of substrates, with an average error of 13 – 19 % [56] and 13 – 23% [57], as well as cellular maintenance coefficients (average error 32 – 41 %) [58], under aerobic and anaerobic conditions. It is noteworthy to point out that these approaches were all developed before inexpensive sequencing technology was readily available to interrogate the details of cellular metabolism, yet they are remarkably predictive [59].

## 4 Genome-scale models of metabolism: peeking in the black box

Due to the advent of low-cost genomic sequencing, and the existence of large databases mapping genes to functions, it has become possible to rapidly characterize the metabolic capability of an organism *in silico*. Consequently, an increasing number of highly detailed genome-scale metabolic models (GEMs) have been developed to systematically understand and engineer microorganisms [60]. If reaction rates *v* are known, the temporal change of metabolite concentrations *x*, is given by 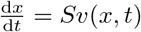, where *S* is the stoichiometric matrix describing the stoichiometry of the metabolic network encoded by the GEM. However, kinetically characterizing each metabolic reaction in an organism is infeasible due to the scale, complexity, and interactions endemic to microbial metabolism, although attempts have been made for two organisms [61, 62]. Consequently, simplifications are typically made to assist analysis. A fundamental assumption made in most GEM studies is that cellular metabolism is approximately in a stationary state. This simplifies the dynamic equation to *Sv* = 0, which relates the stationary reaction fluxes *v* to the system’s stoichiometry alone. Because this central equation is under-determined, further assumptions are necessary. Flux balance analysis (FBA) casts a GEM into a linear program, as shown in Eq. (7).

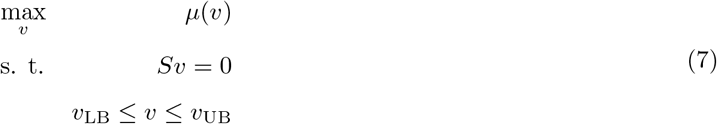

Often, a so-called biomass objective function is used as *µ*; this ad hoc linear sum of weighted fluxes describes the proportion of macromolecular constituents needed by a microorganism to grow [63], and is related to the macrochemical equation used in black box models. Flux measurements and thermodynamic considerations are used to set the flux bounds, *v*_LB_ and *v*_UB_ [64]. It is particularly important to constrain directions of reactions to reduce the flux variability in a model, and to remove thermodynamically infeasible cycles. Typically, physiologically observed metabolite concentration bounds are used in conjunction with Eq. (2) to bound Δ_r_*G*^*′*^, and subsequently assign direction to reactions [65].

While the aforementioned approach ensures that individual reactions obey the laws of thermodynamics, it does not ensure that the entire reaction network is thermodynamically consistent. This can be achieved by imposing additional thermodynamic constraints on the classic FBA algorithm. To this end, the two most robust methods are thermodynamic-FBA [66, 67] and loopless-FBA [68]. The former method directly incorporates thermodynamic parameters, i.e. Δ_r_*G*^*′∘*^, for each reaction, as well as bounds on metabolite concentrations achievable by the cell. By incorporating Eqs. (2) and (3) into (7), the optimization problem is converted into a mixed integer linear program (MILP). The latter method does not require thermodynamic information, however it also reformulates (7) as a MILP. In both cases it is guaranteed that solutions represent thermodynamically consistent flux distributions, albeit at the cost of solving a more computationally intensive optimization problem. Interestingly, while loopless-FBA does not directly use thermodynamic parameters, it can be restated as a simplification of thermodynamic-FBA, in which internal metabolite concentrations are unbounded [69].

Thermodynamic-FBA has been used to investigate the metabolism of *Escherichia coli*. Interestingly, by imposing thermodynamic constraints on its GEM, fewer reactions were found to be directionally constrained than were manually constrained in the original FBA formulation, suggesting that the individual reaction based approach may be too conservative [70]. However, this analysis also highlighted the sensitivity of thermodynamic-FBA to uncertainties in Δ_r_*G*^*′∘*^, as well as metabolite concentration bounds, suggesting that the loopless approach may be more robust, because it does not depend on specific values of the reaction energies. However, computational complexity issues plague both of these MILP-based methods, although faster variants of loopless-FBA have been developed [71]. For this reason, alternative methods that remove thermodynamically infeasible cycles, but do not interrogate reaction directions, are more often used in practice [72, 73].

The rate/yield trade-off can also be quantitatively analyzed with GEMs, if kinetic data is also available. Enzyme flux cost minimization is a technique that combines GEM based analyses with enzyme cost minimization analysis. By using this technique, it was found that environmental, as well as kinetic, parameters affect the trade-off between rate and yield in *E. coli* [74]. The flexibility of constraint based analysis also facilitates its extension, e.g. by incorporating phenomenological observations as constraints on fluxes. Recently, an upper bound on the rate of Gibbs free energy dissipation of cellular metabolism was introduced to further constrain the solution space of FBA [75]. The additional constraints allowed FBA to accurately predict overflow metabolism in *Saccharomyces cerevisiae* and *E. coli*. As argued in [76], a common feature of all models explaining overflow metabolism is the existence of two constraints, which restrict metabolism under different conditions. In [75], these constraints were based on thermodynamic considerations. Interestingly, the same results were obtained by using a far simpler black box approach [77], which suggests that the two approaches could be synergistically combined to yield novel insights.

## 5 Connecting the black box and genome-scale metabolic modeling approaches

Given the increasing ubiquity of GEMs, a rich tool set has been developed to quantitatively analyze metabolism at the genome-scale [78]. However, due to the computational complexity associated with thermodynamic-FBA approaches, it seems that most thermodynamic analyses of GEMs are restricted to reaction direction assignments. In contrast, the historic black box approaches made prodigious use of thermodynamics to overcome their inability to inspect the metabolic features of an organism in detail. Given the accuracy of yield predictions made by using only black box techniques, one can only wonder to what degree deeper thermodynamic integration of GEMs can increase their predictive abilities.

Promisingly, the development of sophisticated thermodynamic databases has already enabled detailed analysis of pathways [41, 43], unavailable a mere decade before their release [79]. These databases can be easily integrated with FBA based approaches. In Fig. 4 we illustrate this integration by calculating the total Gibbs energy dissipated for a variety of organisms and carbon sources, as well as the associated biomass yields.

**Figure 4:**
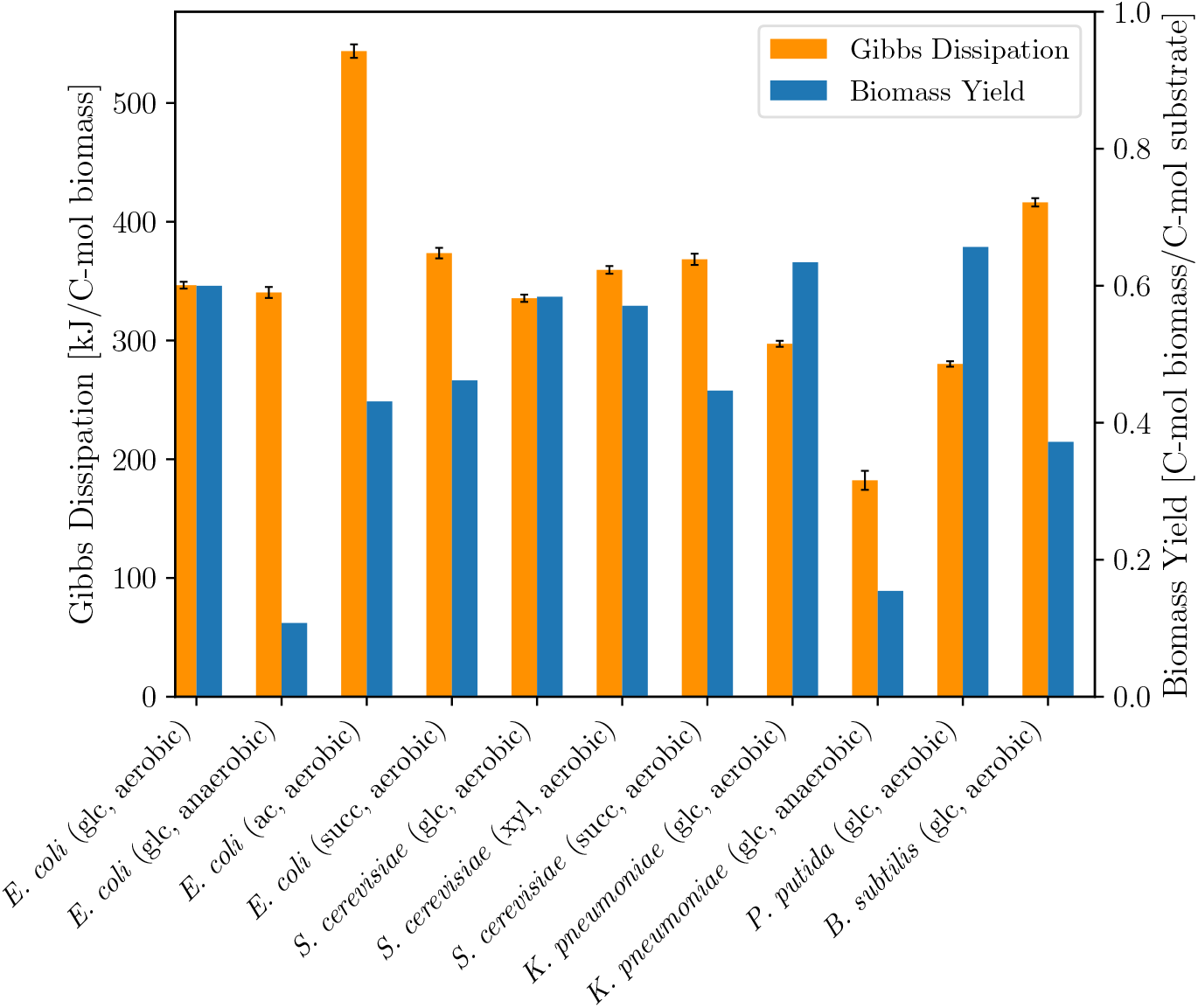
Gibbs dissipation and biomass yields calculated using FBA and the GEMs of *E. coli* [80], *S. cerevisiae* [81], *Klebsiella pneumoniae* [82], *Pseudomonas putida* [83], *and Bacillus subtilis* [84]. Thermodynamic data was sourced from [19] and [85]. Carbon source abbreviations: glc/glucose, ac/acetate, succ/succinate, xyl/xylose. Error bars represent the propagated standard deviation of the mean Gibbs energy of formation for extracellular metabolites and biomass. Code used to generate the plot is available at https://gitlab.com/qtb-hhu/thermodynamic-linear-converter-review-2021.

While it is intuitively clear that catabolic processes drive biomass formation, as modelled in the black box approaches of Section 3, it is less clear how this idea can be applied to GEMs. While GEMs reveal exactly which biochemical transformations are taking place, they do not couple energetics to yield, e.g. there is no disadvantage in increasing the ATP yield of catabolism. However, a rate/yield trade-off must exist. By increasing the number of ATPs generated through catabolism, the Gibbs free energy change of the pathway will approach equilibrium. In light of Sections 2 and 3, this would reduce the rate, since the driving force is diminished. Consequently, metabolism would slow down, reducing the benefit of increasing the ATP yield. This effect is not captured by current constraint-based approaches [86]. It is still an open research question on how to best leverage the bounty of information contained in GEMs, while also incorporating thermodynamic driving force considerations. Addressing this shortcoming by making use of the linear energy converter ideas touched upon in Section 3 could clear the way for establishing the connection between rate and yield at the genome-scale.

## 6 Conflicts of interest, acknowledgements, funding information and author contribution

### Author Contributions

Conceptualization, O.E., S.E.W.; formal analysis, S.E.W., V.V.F, N.P.S., O.E.; investigation, S.E.W., V.V.F., N.P.S., O.E.; writing preparation and review, S.E.W., V.V.F., N.P.S., O.E.; funding acquisition, O.E. All authors have read and agreed to the published version of the manuscript.

## Footnotes

1 In the words of Clausius who originally formulated the second law, “No process is possible whose sole result is the transfer of heat from a body of lower temperature to a body of higher temperature.”

2 Concentrations of all species 1 M, temperature 298.15 K, and pressure 1 bar.

